# PHANES: an integrated resource of protein features for the human proteome

**DOI:** 10.64898/2026.10.14.756639

**Authors:** Dimitrios Miltiadis-Vrachnos, Alexandros Armaos, Laura Broglia, Michele Monti, Jonathan Fiorentino, Gian Gaetano Tartaglia

**Author notes:** To whom correspondence should be sent.

## Abstract

Protein analysis and machine-learning workflows increasingly rely on large-scale numerical representations of proteins. While protein language models provide powerful embeddings, their limited interpretability and high computational cost hinder their use in many biological applications. We present PHANES, a human proteome database and web server providing a comprehensive, biologically interpretable set of features for 20335 proteins. PHANES integrates sequence-derived physicochemical descriptors, structure-based and graph-theoretical features computed from AlphaFold predictions, thermostability indices, and tissue expression specificity. In addition to precomputed features for the human proteome, an integrated web server enables on-demand feature calculation for user-provided protein structures. PHANES supports data export, visualization and comparative analysis at the level of individual proteins and protein groups, bridging the gap between complex numerical representations and biological interpretation while providing an interpretable foundation for exploratory analyses and downstream machine-learning applications. PHANES is freely available at https://tools.tartaglialab.com/phanes.

## 1. INTRODUCTION

Quantitative protein representation via features or embeddings is central to modern computational biology tasks. Recent advances in protein language models (PLMs) have shown that protein embeddings grasp rich structural and functional information (Bepler and Berger 2021, Leclercq and Droit 2025). This enables high quality performance in structure, functionality or custom prediction tasks (Brandes et al. 2022, Elnaggar et al. 2022, Lin et al. 2023). On the downside, these representations cannot be directly interpreted biologically and require substantial computational resources for training or fine-tuning, limiting their applicability in many biological applications.

To address these challenges, we introduce the Proteome-wide Human Atlas of iNterpretable fEatureS (PHANES), a database and web server providing a comprehensive and interpretable ensemble of protein descriptors for the human proteome. PHANES integrates sequence-derived physicochemical properties, structural, graph-based and thermostability features extracted from AlphaFold predicted 3D structures (Jumper et al. 2021) and tissue abundance specificity descriptors using data from PaxDb (Huang et al. 2025). This feature ensemble provides a reusable and reproducible representation of proteins that facilitates biologically interpretable analysis and is ideal for machine-learning workflows.

## 2. IMPLEMENTATION

### 2.1. Data sources and protein set

Predicted protein structures were obtained from the AlphaFold Protein Structure Database (version 6) (Varadi et al. 2021, Bertoni et al. 2026). This resource contains 23586 predicted protein structures for the human proteome. Proteins with sequence length > 2700 amino acids were excluded, as AlphaFold represents these as 1400 aa long, overlapping fragments (Jumper et al. 2021). In the overlapping regions, the same residues can yield different structure-derived feature values depending on the fragment from which they are calculated, preventing an unambiguous assignment of features to the full-length protein. Human proteins over 2700 amino acids were identified using UniProtKB (Swiss-Prot, TrEMBL) (UniProt Consortium 2025), resulting in the exclusion of 216 proteins, which correspond to 3251 Alphafold structures and a final dataset of 20335 proteins.

### 2.2. Feature Categories

PHANES comprises 243 features describing protein sequence, structure, topology, thermostability and tissue specificity context. Features are stored either as scalar values summarizing over the whole protein or as per-residue vectors. A detailed list of all the features is provided in Supplementary Table S1.

#### Sequence-based Features

Physicochemical properties are computed directly from protein sequence. These descriptors are derived from a curated collection of 80 experimentally derived physicochemical propensity scales that capture fundamental biochemical and structural characteristics of proteins (Klus et al. 2014). The scales encode properties related to hydrophobicity, aggregation, membrane association, nucleic acid binding, intrinsic disorder, burial, and secondary structure propensities, including α-helix, β-sheet, and turn formation, as well as two phenomenological sequence patterns (Klus et al. 2014). Such physicochemical properties provide mechanistically interpretable descriptors of sequence-dependent molecular behaviour and have been implicated in processes ranging from molecular recognition to multivalent interactions and phase separation (Fiorentino, Monti, Broglia et al. 2026). For each descriptor, per-residue values and protein-level averages are provided. In this category, a value for each amino acid is assigned, which is averaged over a sliding window of 7. These descriptors have been previously implemented in tools such as cleverSuite (Klus et al. 2014), employed in the training of the catGRANULE 2.0 algorithm (Monti et al. 2025) for predicting liquid–liquid phase separation (LLPS) propensity (Fiorentino et al. 2026) and the HERCULES algorithm for predicting RNA-binding propensity, and in theoretical analyses on the informativeness of protein features (Fiorentino et al. 2026), demonstrating their robustness and predictive relevance.

#### Structure-based Features

Structure-based descriptors are derived from AlphaFold2-predicted models downloaded from the AlphaFold Protein Structure Database (Jumper et al. 2021, Varadi et al. 2021, Bertoni et al. 2026). Prediction confidence is summarized using the mean and standard deviation of per-residue predicted Local Distance Difference Test (pLDDT) scores. Structural features are computed from PDB files using the Bio.PDB module of Biopython (v1.81). Extracted descriptors include the total number of residue–residue contacts and contacts normalized by protein length, as well as the radius of gyration (Rg) and length-normalized Rg to quantify structural compactness. Solvent accessibility is assessed by computing the relative Accessible Surface Area (ASA) of the amino acid which is the fraction of the exposed area of the amino acid divided by its total surface area. In this manner, residues are classified as internal (ASA < 0.5) or exposed (ASA ≥ 0.5). For both internal and exposed residues, we compute secondary structure content (helix, strand, coil, turn, disorder), accessibility, and charge distribution. Together, these features provide a compact yet comprehensive description of protein packing, compactness, solvent exposure, electrostatic organization, and structural confidence based on AlphaFold predictions.

#### RNA-binding patch Features

To characterize RNA-binding propensity, PHANES includes 9 descriptors capturing compositional similarity between regions of the protein and experimentally determined RNA-binding domains (RBDs) (Castello et al. 2016, Monti et al. 2025). A sliding window is applied along the protein sequence and for each window the amino acid composition is computed and compared to the composition of peptides derived from a curated set of experimental RBDs (Castello et al. 2016). For each region, the compositional distances to all reference RBDs are calculated, and the distances and indices of the three most similar domains are retained as features. This procedure is performed separately for internal residues, exposed residues, and the full-length protein sequence. These descriptors provide a structure-aware measure of similarity to known RNA-binding regions, integrating sequence composition with residue exposure information.

#### Graph Network theoretical features

In this category, proteins are represented as Residue Interaction Networks (RINs) following a modified graph-based framework inspired by (Miotto et al. 2019). In these networks, nodes correspond to amino acids and edges represent residue–residue interactions, defined when the spatial distance between residues is ≤ 12 Å. The networks are implemented using the NetworkX Python library. From each RIN, 31 graph-theoretical descriptors are extracted and grouped into four categories: general graph, centrality, hub, and clustering features. General graph metrics include global properties such as the number of nodes and edges. Centrality features capture the relative importance of residues within the network and include measures such as Katz and closeness centrality. Hub-related descriptors include hub and authority scores, identifying residues with prominent connectivity patterns. Clustering features quantify local cohesiveness and modular organization, including metrics such as clustering coefficient and triangle counts. Most graph-based descriptors are computed at the residue level and stored as per-residue vectors, alongside their corresponding protein-level summary statistics (e.g., mean values). This dual representation captures both local interaction patterns and global topological organization, providing a comprehensive network-based characterization of protein structure.

#### Thermostability features

Thermostability-related descriptors are computed using the Thermometer framework (Miotto et al. 2019, 2022), which estimates protein thermal stability through a graph-based energetic model. A total of nine features are included, derived from three Thermometer stability (Ts) descriptors corresponding to total interaction energy, Lennard–Jones interactions, and Coulombic energy. Each energy term is calculated separately for polar, non-polar, and combined residue interactions. These descriptors quantify the deviation in interaction energies between the native protein structure and an ensemble of randomized residue interaction networks. The randomized networks preserve the original nodes and overall connectivity architecture, while edge weights are reassigned by sampling from energy distributions characteristic of mesostable proteins. The resulting Ts scores therefore quantify how strongly the energetic organization of a protein deviates from the mesostable reference distribution. Values showing a larger deviation toward the thermostable region indicate an interaction network more characteristic of thermostable proteins, whereas values closer to the mesostable reference indicate a more mesostable-like energetic organization. The three energy-specific Ts descriptors are intended to be interpreted jointly rather than as direct estimates of melting temperature.

#### Tissue specificity features

Two expression-based descriptors capture protein abundance and tissue specificity using PaxDB data across 63 human tissues (Huang et al. 2025). One descriptor summarizes the overall abundance of a protein across tissues, whereas the second quantifies how restricted its expression is to a subset of tissues. Together, these features distinguish broadly expressed proteins from tissue-specific proteins and provide biological context complementary to sequence- and structure-based descriptors.

### 2.3. Web Server and user input

The PHANES web server enables interactive querying by UniProt accession and supports feature selection, filtering and export in csv/pkl format. Users may also upload up to ten protein structures (PDB files) to compute PHANES features on demand via an integrated web server. For non-human or synthetic proteins, expression-based features are omitted. Thermometer calculations are applied for proteins less than 1000 amino acids long, extending the original threshold of the application.

## 3. Results

PHANES provides an integrated environment for feature extraction, visualization, and comparative analysis at both the single-protein and protein-group levels. Proteins are queried using UniProt accession numbers, followed by flexible selection and filtering of feature categories (Fig 1A). The resulting structured dataset, containing the selected proteins and descriptors, can be downloaded in csv format for downstream analyses. In parallel, the platform offers a diverse suite of visualization tools to facilitate exploratory data analysis directly within the interface.

**Figure 1.**
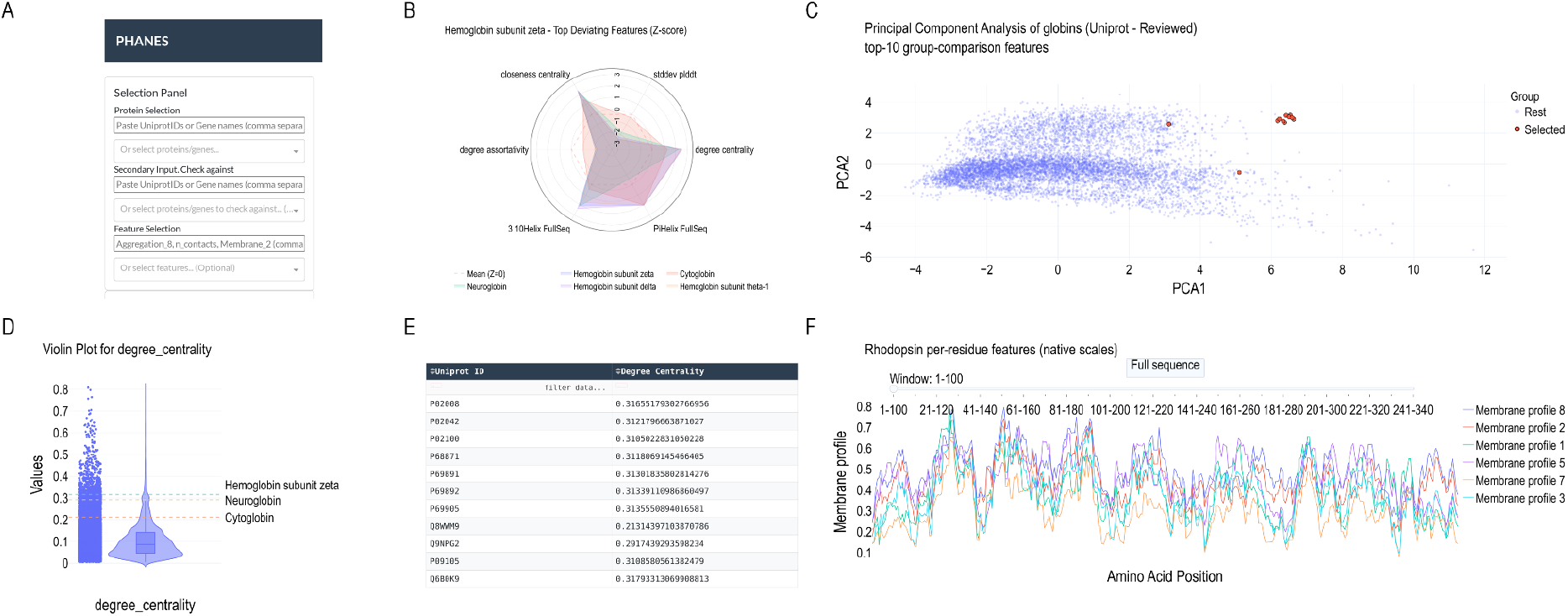
The PHANES application and representative analysis modules. (A) Selection Panel. Proteins are queried by Uniprot ID or gene names; an optional second set can be supplied to compare against, and the features to analyse are chosen similarly. (B) Spider Plot. Per-protein z-scores of the features that deviate most from the whole-proteome mean, shown for selected globins (P02008, P02042, Q8WWM9, Q9NPG2). Each axis is one top-deviating feature; the dashed contour marks the proteome mean (z = 0), so distance from the center reflects the magnitude and direction of the deviation. (C) Principal Component Analysis (PCA). Proteome-wide PCA computed on user-selected features that mark the globin set (red marks) against the human proteome (blue marks). (D) Violin Plot. Distribution of a single selected feature across the whole proteome; dashed lines indicate values of selected proteins, locating them within the population. (E) Feature matrix. Tabular per-protein values of the selected features for the according queried proteins; the table is sortable, filterable and can be exported. (F) Per-residue Plot. Position-resolved values of selected per-residue features (Membrane_1,*2,3,5,7*,8) along the rhodopsin (P08100) sequence, shown on their native scales. A sliding window and a “Full sequence” button control and allow inspection of local regions or the whole protein.

### Single-Protein Analysis

For individual proteins, PHANES enables detailed characterization through spider plots, per-residue feature profiles, and distribution-based visualizations such as violin plots. Spider plots highlight the features for which a selected protein deviates most strongly—based on z-scores—from a chosen reference set, such as the human proteome or a user-defined protein group (Fig 1B). This representation allows rapid identification of distinctive properties, shared trends, or potential inclusion within functional groups. Violin plots provide distributional context for scalar features, illustrating spread, skewness, and locality relative to the reference population (Fig 1D). The position of the queried protein within this distribution allows users to assess whether a given property is typical or unusually high or low compared with the human proteome, providing context for interpreting individual feature values.. Per-residue line plots leverage vector-based descriptors stored in the database and can be explored in two complementary ways: (i) visualizing multiple features along a single protein sequence to identify consensus regions or structurally/functionally enriched segments, and (ii) visualizing a single feature across multiple proteins to detect conserved motifs or region-specific differences (Fig 1F). Together, these tools enable both global and region-resolved protein characterization.

### Protein-Group Analysis

A central application of PHANES is the systematic comparison of proteins and protein groups to identify shared characteristics and properties that distinguish different protein families or functional classes. Users can compare predefined sets of proteins or investigate how selected proteins differ from the human proteome as a whole. By examining multiple interpretable descriptors simultaneously, these comparisons can reveal whether similarities or differences are primarily associated with sequence composition, structural organization, thermostability, or tissue specificity. Dimensionality reduction techniques — including Principal Component Analysis (PCA) and Uniform Manifold Approximation and Projection (UMAP) — enable clustering identification and visualization in reduced feature space (Fig 1C). These methods can be applied to the full feature set or to user-defined subsets tailored to specific biological questions. Heatmaps provide an overview of inter- and intra-group similarities, highlighting feature-level patterns and divergences across proteins. Additionally, protein similarity networks can be constructed, where nodes represent proteins and edges connect pairs with cosine similarity above a user-defined threshold. This graph-based representation offers an alternative clustering strategy and allows flexible adjustment of similarity criteria. Finally, PHANES identifies features that discriminate between two protein groups—or between a user-defined group and the human proteome—allowing users to investigate which biological properties contribute to the observed differences. This functionality supports hypothesis generation, comparative characterization of protein families, and the construction of interpretable datasets for downstream predictive modeling.

### Data & Features

The proteins and features selected by the application’s inputs are displayed in a matrix with the original data (Fig1E). This makes user specified data downloadable on demand in csv format. A spearman’s ρ (Spearman 2010) correlation heatmap is provided for illustrating pairwise feature relationships in order to provide insights for feature selection pipelines.

## 4. Availability

PHANES, along with documentation, tutorials and feature description, is freely available at https://tools.tartaglialab.com/phanes. The web server is compatible with modern browsers.

Source code is available at [https://github.com/tartaglialabIIT/phanes].

## Funding

The research leading to these results have been supported through ERC [ASTRA 855923, H2020 Projects IASIS 727658 and INFORE 825080 and IVBM4PAP 101098989 and National Center for Gene Therapy and Drug based on RNA Technology (CN00000041), financed by NextGenerationEU PNRR MUR - M4C2 - Action 1.4- Call “Potenziamento strutture di ricerca e di campioni nazionali di R&S” (CUPJ33C22001130001).

## Acknowledgments

The authors acknowledge the members of the RNA Systems Biology Lab at IIT for useful discussions.

## Notes

https://tools.tartaglialab.com/phanes

https://github.com/tartaglialabIIT/phanes.git

### Competing Interest Statement

The authors have declared no competing interest.

